# Intrastriatal delivery of a zinc finger protein targeting the mutant HTT gene by AAV9 obviates lipid phenotypes in brain and plasma of zQ175DN HD mice

**DOI:** 10.1101/2025.01.06.630918

**Authors:** Andrew Iwanowicz, Adel Boudi, Connor Seeley, Ellen Sapp, Rachael Miller, Sophia Liu, Kathy Chase, Kai Shing, Ana Rita Batista, Miguel Siena-Esteves, Neil Aronin, Marian DiFiglia, Kimberly B. Kegel-Gleason

## Abstract

Reducing the burden of mutant Huntingtin (mHTT) protein in brain cells is a strategy for treating Huntington’s disease (HD). However, it is still unclear what pathological changes can be reproducibly reversed by mHTT lowering and whether these changes can be measured in peripheral biofluids. We previously found that lipid changes that occur in brain with HD progression could be prevented by attenuating HTT transcription of the mutant allele in a genetic mouse model (LacQ140) with inducible whole body lowering. Here, we tested whether intrastriatal injection of a therapeutic capable of repressing the mutant *HTT* allele with expanded CAG can provide similar protection against lipid changes in HD mice with a deletion of neo cassette (zQ175DN). Wild-type or zQ175DN mice were injected with AAV9 bearing a cDNA for a zinc finger protein (ZFP) which preferentially targets mutant HTT (ZFP-HTT) to repress transcription (1). Proteins from brain tissues were analyzed using western blot, capillary electrophoresis, and nitrocellulose filtration methods. Lipid analyses of brain tissue and plasma collected from the same mice were conducted by liquid chromatography and mass spectrometry (LC-MS). Somatic instability (SI) index was assessed using capillary gel electrophoresis of PCR products and was shown to be impeded by HTT-ZFP. Lowering mHTT levels by 43% for 4 months prevented loss of total lipid content including subclasses sphingomyelin (SM), ceramide, phosphatidylethanolamine (PE) and others of caudate-putamen in zQ175DN mice. Moreover, LC-MS analysis of plasma demonstrated total lipid increases and lipid changes in monogalactosyl monoacylglyceride (MGMG) and certain phosphatidylcholine (PC) species were reversed with the therapy. In summary, our data demonstrate that analyzing lipid signatures of brain tissue and peripheral biofluids are valuable approaches for evaluating potential therapies in a preclinical model of HD.

**Funding:** CHDI Foundation, Dake Family Fund

**Disclosure statement:** The authors have nothing to disclose.

**Author contributions:** AI and KS extracted lipids and performed computational and statistical analysis; AB and CS collected plasma and brain tissues; SL and ES performed protein chemistry; KC maintained mouse colonies, RM performed stereotaxic injections, RB subcloned ZFP cDNAs and prepared virus, MSE, NA, MD, and KKG planned experiments and wrote manuscript.

## Introduction

A heritable mutation causing abnormal expansion of a normal CAG repeat sequence in the *HTT* gene causes Huntington’s disease (HD) (2), a neurodegenerative disease in which patients suffer progressive cognitive, psychiatric, and motor symptoms, and ultimately death (3). Further CAG repeat expansion can happen in individual cells with age (somatic instability, SI), and genes involved in DNA mismatch repair were identified as modifiers of age of onset in human GWAS studies (4, 5).

Because mutant *HTT* gene products may be involved in diverse pathological mechanisms, gene therapy approaches to block the expression of *HTT* gene products may be superior to other therapies targeting downstream mechanisms. One approach is to target the *HTT* gene directly at the level of transcription using proteins engineered to bind the mutant allele, such as a zinc finger protein (ZFP) which has been shown to selectively repress transcription from the mutant HTT allele containing high CAG repeats (1). This ZFP fusion protein has the advantage of blocking all gene products, including the small HTT1a exon1-intron1 readthrough transcript (1).

Many of the experimental readouts reported in studies using HD mouse models are semi-quantitative, have high variability requiring numerous mice and are labor intensive. The readouts used to assess the effects of HD gene lowering include mRNA levels for striatal genes known to change in HD (PDE10 and DARPP32) (6), immunofluorescence for DARPP32, ligand binding to PDE10, dopamine receptors 1 and 2 (1), brain mass, volume of caudate-putamen, the presence of inclusions formed by mHTT, and behavioral and cognitive measures all assessed at one or two time points post treatment (7-9).

RNAseq or microarray analysis are quantitative and give strong, reproducible phenotypes in HD mouse models (10, 11) and have become an essential quantitative readout for lowering therapies.However, there was limited improvement in reversing changes in levels of mRNA transcripts specific to striatal medium spiny neurons (MSN) in brains treated with an ASO targeting *HTT* (6). And only partial reversal of the transcriptomic signature occurred with a genetic inducible model of mHTT lowering which lowered in all cell types (LacQ140; (12)).

Using the highly accurate and quantitative method of liquid chromatography and mass spectrometry (LC-MS), we recently demonstrated in the LacQ140 mice that significant changes in levels of numerous lipids occurred, and reducing mHTT levels was protective (13). Here, we tested whether the benefits of an injected therapeutic could be evaluated using LC-MS to quantitatively track global lipid changes in brain and in a peripheral biofluid in the Q175 HD mouse model.

## Methods and Materials

### Animal Welfare Statement

All the studies in this manuscript were conducted in compliance with the Institutional Care and Use Committee (IACUC) guidelines of the University of Massachusetts Chan Medical School (docket #202100018).

### AAV subcloning & packaging

Details of the AAV subcloning and packaging can be found in **Figure 1A and Supplemental Methods**.

**Figure 1.**
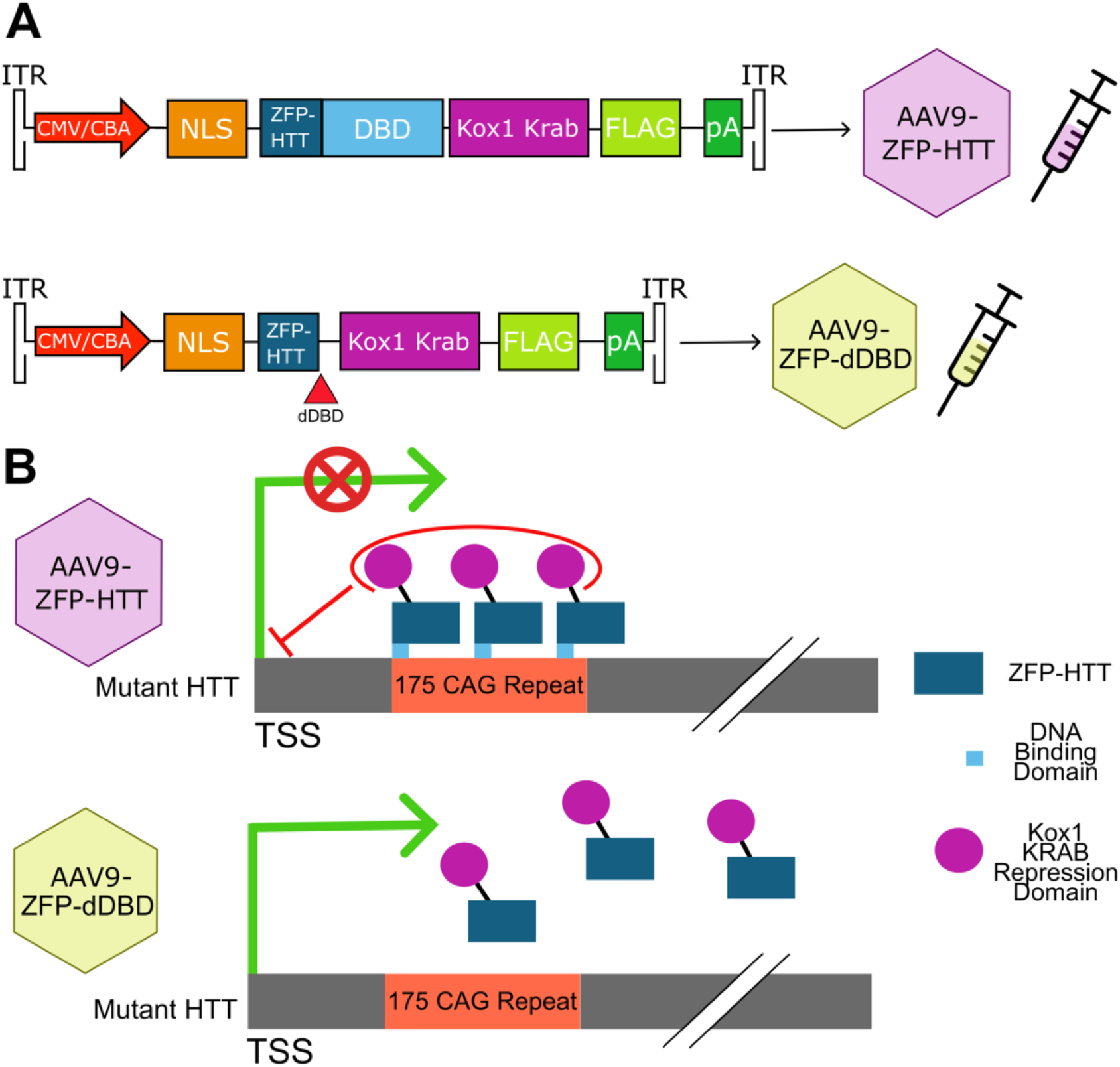
Diagram of AAV9-ZFP-HTT and AAV9-ZFP-dDBD constructs. **(A)** Diagram of the cDNA encoding construct “30645” specific to mutant HTT (ZFP-HTT) was fused in frame with the KRAB repressor domain (Construct D from Zeitler B. Nature Med 2019) and subcloned into the multiple cloning site of the self-complimentary adeno-associated virus 9 (AAV9) vector behind a chick beta-actin promotor. This fusion protein has been shown to selectively repress transcription of the mutant HTT allele containing high CAG repeats including the HTT1a exon1-intron1 readthrough transcript (1).The DNA-binding domain (DBD), amino acids 16-170 out of 278, was removed in the dDBD construct. **(B)** Schematic mechanism of action of the ZFP transcriptional with and without the DNA binding domain.

### Mouse Intracranial Injection Surgeries and tissue collection

Heterozygous zQ175DN HD mice (abbreviated to HD in figures and tables) and their wild-type (WT) littermate controls (Q7/Q7) were purchased from The Jackson Laboratories (Z_Q175 KI (neo-); (C57BL/6J)-CHDI stock # 370832, JAX stock #029928). Briefly bilateral intrastriatal injections of AAV9 virus (3.5 × 10^10^ PFU) were performed at eight weeks of age. At 6 months of age brain tissue and plasma was collected. More details can be found in **Supplemental Methods**.

### Tissue fractionation, SDS-PAGE, western blot analysis, capillary gel electrophoresis and filter trap assay

Details can be found in **Supplemental Methods**.

### Lipid Extraction, LC-MS, and Annotations

Lipids were extracted using the two-phase methyl-*tert*-butyl ether (MTBE) method as previously described (14). Details in **Supplemental Methods**.

### Computational analysis/visualization

Pixel intensity quantification for immunoblotting was performed using ImageJ, more details can be found in **Supplemental Methods**. To correct for duplicate annotation of lipid ions by LipidSearch caused by wide elution profiles, all instance of identical lipid ions were summed together. Lipid ions were then filtered using the LipidSearch parameter (Rej. = 0) and lipid ions missing in over 20% of the samples were excluded from analysis. Data was then log-transformed and normalized with Eigen MS (15). The principal component analysis was performed using FactoMineR (v2.11), data was centered and scaled. Total level of lipid in each sample was determined by summing the intensity of all the lipid ions in each sample. The heatmap in figures five and six were generated in R using ComplexHeatmaps (v2.20.0; (16)). Statistical significance of lipid subclasses was determined by two-way ANOVA between all three groups and sex, followed by Tukey’s t-test (p value), and the false discovery rate (FDR, 5%) (q value) was controlled by the two-stage linear step-up procedure of Benjamini, Krieger, and Yekutieli (brain lipid subclass N=42, brain lipid ions N=2398, plasma lipid subclass N = 47, plasma lipid ions N=2251). A lipid subclass/ion was considered recovered if the sign of the log_2_-Fold change was the same when comparing WT treated with ZFP-dDBD and HD mice treated with ZFP-HTT to HD mice treated with ZFP-dDBD, and the q value between the HD treatment groups was less than 0.05.

### Somatic Instability index

Details can be found in **Supplemental Methods**.

## Results

### Lowering of mutant HTT protein in Q175DN HD mice

To test whether a gene therapy capable of allele-specific gene repression could prevent lipid-related neurodegenerative changes in an HD preclinical mouse model, prepared virus for AAV9-ZFP-HTT or AAV9-ZFP-dDBD (**Fig. 1B**) were introduced into the zQ175DN HD mouse caudate-putamen with bilateral injections at two months of age, prior to known behavioral phenotypes in this model (17, 18). Wild-type (WT) Q7/Q7 mice were injected with AAV9-ZFP-dDBD virions as an additional control to obtain baseline levels of genotype-dependent changes in proteins and lipids. All tissues were harvested at six months of age when considerable changes in transcription have been described in zQ175/Q7 striatum compared to WT, nuclear and cytoplasmic aggregates are present and electrophysiological changes can be detected (1, 10).

To examine HTT protein levels, we performed capillary gel electrophoresis on crude homogenates of AAV-treated caudate-putamen of HD mice using anti-polyglutamine antibody, MW1, to detect mutant HTT protein (mHTT). Results showed a significant lowering (48%) of mHTT with expression of ZFP-HTT but not with ZFP-dDBD in caudate-putamen of HD mice (**Fig. 2A, B**). We also performed western blot (WB) assays using the crude homogenates from both WT and HD treated mice and probed with anti-N-terminal HTT antibody Ab1, which detects both WT HTT and mHTT (**Fig. 2C, D**, (13, 19, 20)). Using this method, pixel intensity quantification showed that the level of mHTT was 43% lower in HD mice treated with ZFP-HTT compared to those treated with ZFP-dDBD (**Fig. 2D**). The levels of WT HTT in heterozygous zQ175DN HD were an average of 54 and 64% lower than in WT mice which have two WT alleles, as expected (**Fig. 2D**). There was no difference in WT HTT levels between the HD treatment groups demonstrating the specificity of the ZFP-HTT repressor protein for the mutant HTT allele. Thus, four months after injection, mHTT in crude homogenates was significantly lowered with ZFP-HTT treatment compared to the control virus (HTT-dDBD) and WT HTT levels were spared.

**Figure 2.**
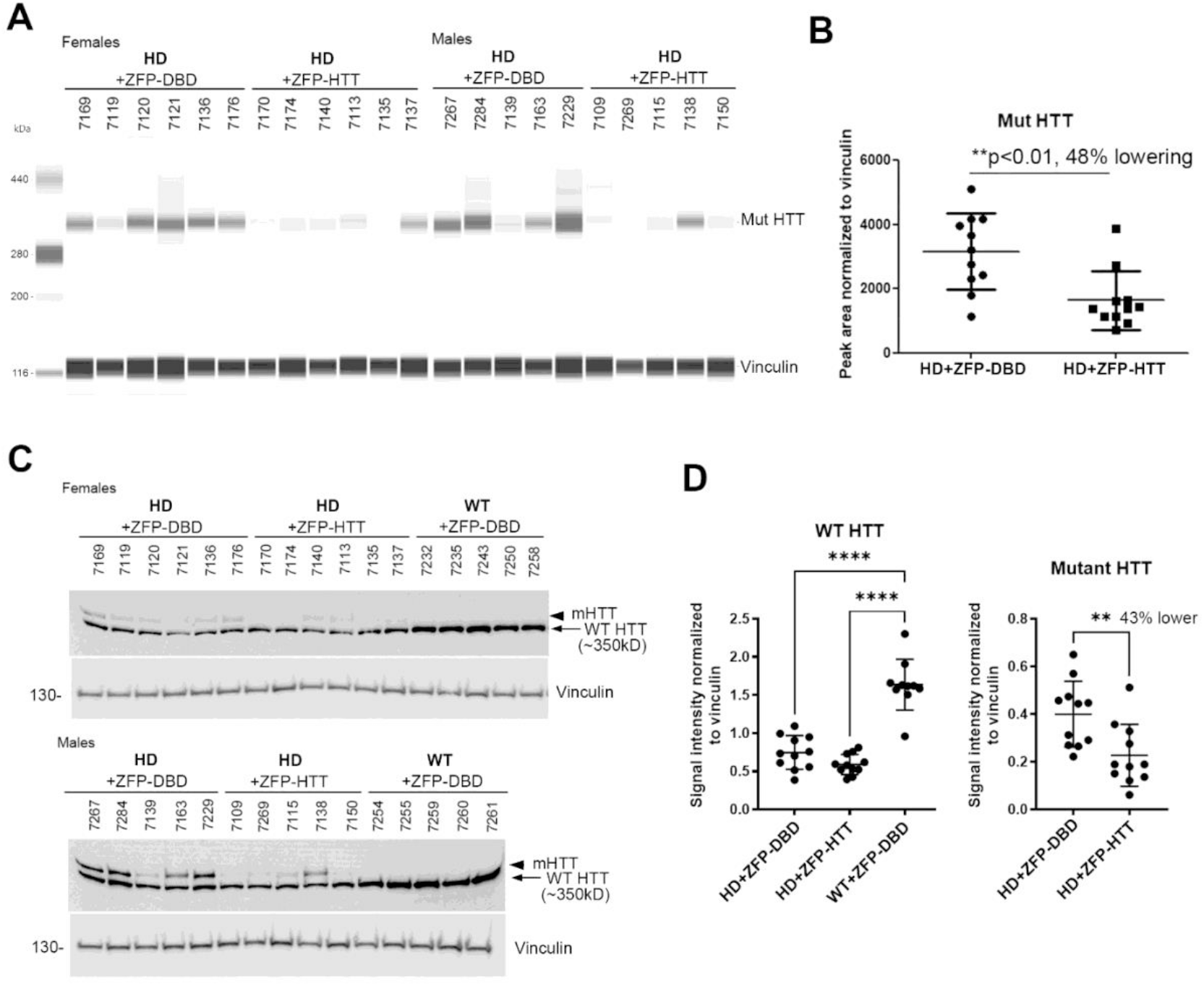
Capillary electrophoresis (WES) analysis and western blot analyses of protein levels of soluble normal and mutant Huntingtin in crude homogenates from zQ175 caudate-putamen injected with AAV9-ZFP-HTT or AAV9-ZFP-dDBD and WT injected with AAV9-ZFP-dDBD. **A**. Lane view image of automated simple western (Wes) using equal amounts of protein (0.6ug) and anti-polyQ antibody MW1. **B**. Quantification of capillary electrophoresis shows significant lowering of mutant HTT by 48% with ZFP-HTT treatment compared to treatment with ZFP-DBD in Q175/Q7 HD mice. **C**. Western blot images show equal protein loading (10 ug) of crude homogenates from female (top) and male (bottom) mice probed with antibodies indicated to the right of the blots. **D**. Graphs show signal intensity normalized to vinculin of the combined female and male samples. There is significantly less WT HTT in the Q175/Q7 HD mice compared to WT mice, as expected (50-60% less, ^****^p<0.0001, One-way ANOVA with Tukey’s multiple comparison test, n=10-11 per group). There is 43% lowering of mutant HTT in the ZFP-HTT treated Q175 HD mice compared to the ZFP-DBD treated HD mice (^**^p<0.01, unpaired t test, n=11 per group).

### A molecular species of mHTT persists with lowering by ZFP-HTT

Previously, we showed that a molecular species of mHTT in the crude nuclear pellet fraction (P1) was resistant to lowering in LacQ140 mice (13). We and others have shown that the anti-HTT antibody S830 detects a slowly migrating species of mHTT that appears as a high molecular weight smear in homogenates from brain tissues containing mHTT (13, 21, 22). Using Ab1 on WB, levels of mHTT were 54% lower in P1 from zQ175DN mice treated with ZFP-HTT compared to ZFP-dDBD (**Fig. 3A, B**). WT HTT levels in the P1 fraction were an average of 50-53% lower in the zQ175DN mice compared to Q7/Q7, as expected for one allele versus two alleles of normal HTT, consistent with findings above in crude homogenates. However, whereas antibody S830 recognized full-length soluble mHTT detected as a resolvable band which was 44% lower in the AAV9-ZFP-HTT mice compared to those treated with AAV9-ZFP-dDBD, no lowering of the high molecular weight smear occurred (**Fig. 3A, C**). This result is consistent with our findings in LacQ140 striatum where a S830-positive species of HTT persists even with transcriptional lowering. Next, a filter trap assay to analyze lowering of aggregated mHTT in P1 was employed. MW8 antibody, which detects aggregated exon1 HTT (21), showed a significant reduction of aggregated mHTT in the zQ175DN mice treated with AAV9-ZFP-HTT compared to those treated with AAV9-ZFP-dDBD (**Fig. 3D**).

**Figure 3.**
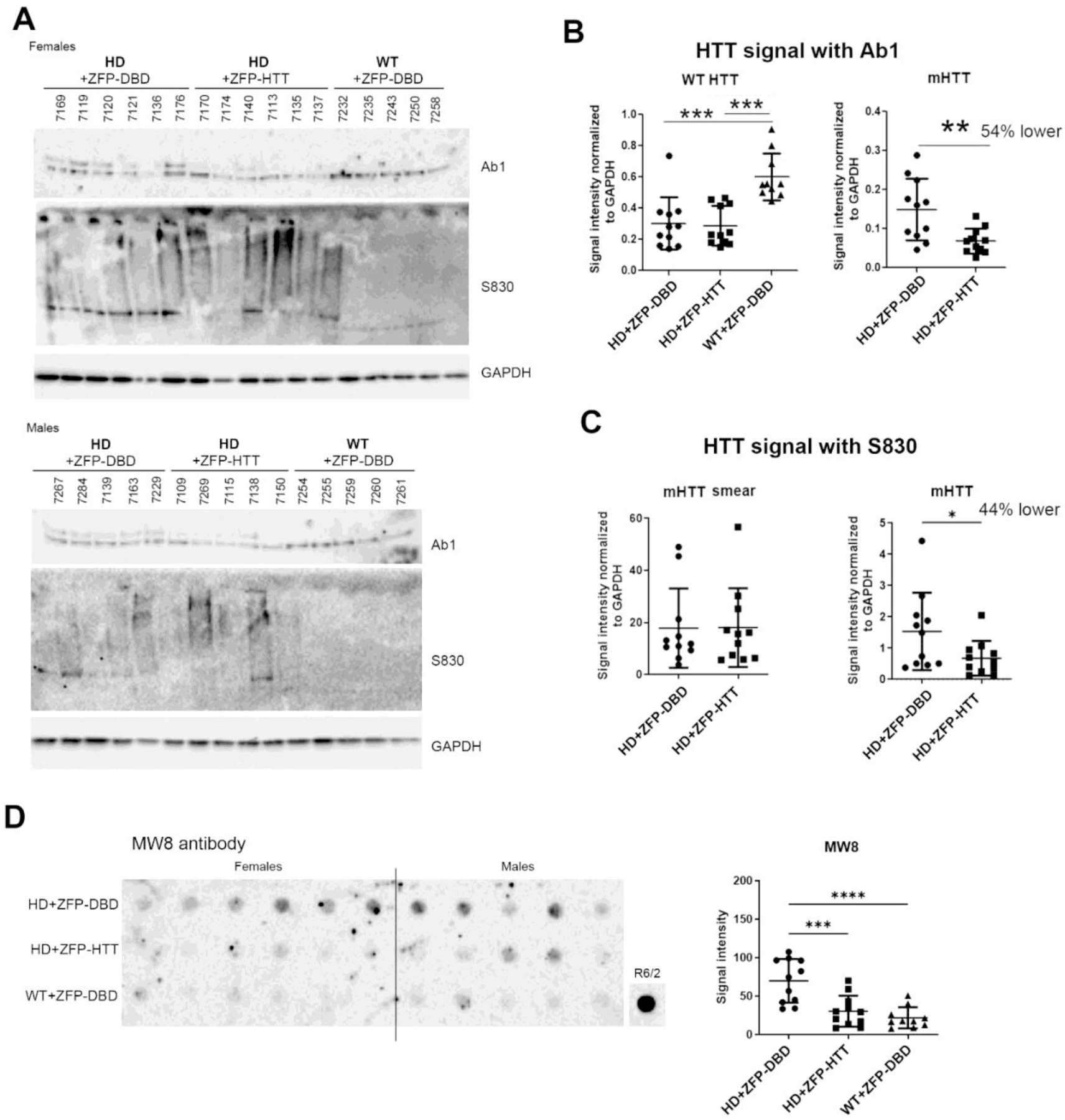
Protein analysis of the crude nuclear P1 fractions from caudate-putamen from mice injected with AAV9-ZFP-HTT or AAV9-ZFP-dDBD and WT injected with AAV9-ZFP-dDBD analyzed by western blot and gel filtration. **A**. Equal amounts of protein (10ug) from the P1 fractions were separated by SDS-PAGE and probed with anti-HTT antibody S830 to detect SDS-resistant high molecular weight HTT protein. Another gel was run with the same samples and probed with anti-HTT antibody Ab1 to detect full-length soluble HTT. **B, C**. No significant difference aggregated HTT in the smear at the top of the gel with S830 was detected in the Q175/Q7 mice treated with ZFP-DBD or ZFP-HTT but there was significant lowering of full-length HTT as detected with Ab1 (54% lower, **p=0.0056, unpaired t test) and S830 (44% lower, *p=0.0494, unpaired t test) in the P1 fraction similar to we saw in the crude homogenates. **D**. Filter trap assay using 40ug samples of the P1 fraction with MW8 antibody shows significantly more aggregated HTT protein on the membrane in HD+ZFP-DBD mice compared to both HD+ZFP-HTT and WT+ZFP-DBD mice and no difference between HD+ZFP-HTT and WT+ZFP-DBD mice (One-way ANOVA with Tukey’s multiple comparison test, ^***^p<0.001, ^****^p<0.0001).

All together, these data indicate that reducing mutant *HTT* transcription with AAV9-ZFP-HTT lowers soluble and aggregated forms of mHTT in crude homogenate and P1 fraction, but that a S830-positive misfolded molecular species of mHTT which is present in the crude nuclear fraction resists lowering.

### AAV9-ZFP-HTT prevents genotype-dependent lipid changes in Q175DN HD caudate-putamen

Previously, we found that age and genotype dependent lipid changes occur in both zQ175/Q7 mice (23) and LacQ140 mice, and these changes could be prevented by early whole body genetic downregulation of mutant *HTT* transcription in LacQ140 mice (13). To test whether intrastriatal injection of a therapeutic would have the same protective effects, we surveyed lipids in crude homogenates of our AAV-treated mice by liquid chromatography and mass spectrometry (LC-MS). LC-MS analysis of lipids from caudate-putamen of treated WT and zQ175DN mice at six months of age yielded high coverage of lipids species with 2,398 unique lipid ions identified across 42 lipid subclasses. To assess the variation within the data, a principal component analysis (PCA) was performed. The projection of the two most informative principal components, PC1 and PC2 are shown in **Figure 4A** (PC1, 14.3% of variance; PC2, 13.0% of variance). Most of the variation along the first principal component was within each group.

**Figure 4:**
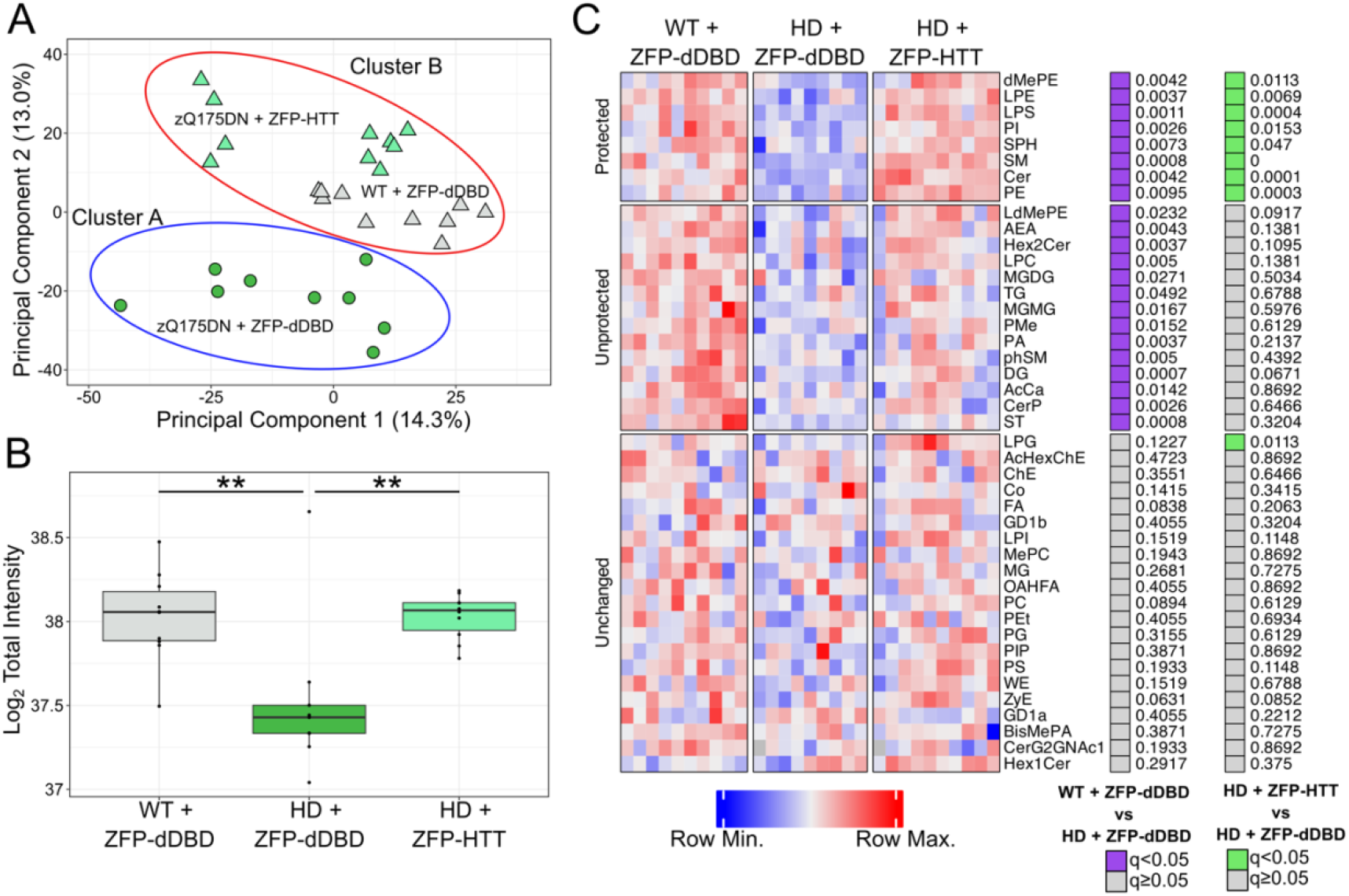
Protection of total lipid levels and subclass levels via lowering of mutant HTT in the caudate-putamen. **(A)** Unsupervised classification methods showed prevention of lipid defects by lowering of mHTT. The first principal component captured 14.3% of variation and showed within group variation. The second principal component captures 13.0% of variation and showed WT mice treated with ZFP-dDBD (white triangles) cluster closer together with HD mice treated with ZFP-HTT (light green triangles) than HD mice treated with ZFP-dDBD (dark green circles). *K-means clustering* (k=2, intial points = 25) on the first eight principal components clustered the HD mice treated with ZFP-HTT with the WT mice treated with ZFP-dDBD together and the HD mice treated with ZFP-dDBD separately. **(B)** Loss of total lipid intensity is prevented by lowering of mutant HTT. Two way Anova revealed signicant differences in treatment (F(2,23) = 8.645; p = 0.00158) but not sex or the interaction between sex and treatment. zQ175DN mice treated with ZFP-dDBD had 30% lower levels of total lipids than WT mice treated with ZFP-dDBD (p=0.0035; **) and HD mice treated with ZFP-HTT (p = 0.0042; **). **(C)** Lowering of mHTT prevents changes in several lipid subclasses. The heatmap depicts individual animals (columns) and lipid subclass intensities (rows). *Purple* boxes indicate significantly different levels between WT + ZFP-dDBD and HD + ZFP-dDBD. *Green* boxes indicate significantly different levels between HD + ZFP-HTT and HD + ZFP-dDBD. (Two-way Anova with Tukey’s multiple comparison test two-stage linear step-up procedure of Benjamini, Krieger and Yekutieli, N=42 lipid subclasses, n=10 mice).

Along the second principal component, all groups showed separation from each other. The HD mice treated with ZFP-HTT are closer to the WT mice treated with ZFP-dDBD. To confirm the PCA results obtained using two components, we undertook a more in-depth quantitative analysis called k-means computational clustering, which considers changes in all lipid levels (increased or decreased). We queried which two of the groups (k=2) were most similar using this method with 25 initial starting points employed on the first eight principal components which accounted for 57% of the total variation. HD mice treated with the control virus clustered into group A (**Fig. 4A, *blue ellipse***) and WT mice treated with the ZFP-dDBD and HD mice treated with ZFP-HTT clustered into group B (**Fig. 4A, *red ellipse***).

This k-cluster analysis showed that HD mice that had received the therapeutic were more similar to WT mice than HD mice that had received the control virus.

An examination of total lipid levels showed a genotype-dependent loss of lipids in HD mouse caudate-putamen (30%) compared to WT mice, each treated with ZFP-dDBD (**Fig. 4B**), consistent with previous results in LacQ140 mice. Changes in several lipid subclasses were prevented by treatment with AAV9-ZFP-HTT. There were 23 subclasses of lipids that were significantly lower in HD mice compared to WT mice treated with ZFP-dDBD (**Fig. 4C and Supplementary Fig. 1**). Levels of eight subclasses were corrected with ZFP-HTT treatment (**Fig. 4C**).

Next we looked at individual lipid species which can have precise functions in signaling such as arachidonic acid signaling (24) and myelin stability (24, 25). ZFP-HTT treatment prevented most genotype-dependent changes in individual lipid species (**Supplementary file 1**). When compared to WT mice treated with the control virus, the HD mice treated with the control virus had 270 significantly altered individual lipid ions. Of those, 58% (157/270) were corrected by the ZFP-HTT treatment **(Table 1)**.

**Table 1.** Lipid Ion changes in HD Caudate-Putamen and Plasma

|  | Number of Annotated Lipid Ions | Number of Lipid Ions altered between WT and HD treated with ZFP-dDBD | Number of Lipid Ions Recovered |
| --- | --- | --- | --- |
| Caudate-Putamen | 2398 | 270 | 157 |
| Plasma | 2251 | 70 | 39 |

### AAV9-ZFP-HTT prevents genotype-dependent lipid changes in Q175DN HD plasma

Lipid alterations have been detected in plasma of HD patients, especially in the phosphatidylcholines and free fatty acids (26-28). To determine if lipid levels in plasma could be an effective peripheral readout in a HD preclinical model, plasma was collected from the same mice used for caudate-putamen studies and analyzed by LC-MS. LC-MS analysis offered excellent coverage of plasma lipids with 2251 lipid ions across 47 subclasses.

There was a genotype-dependent increase in total lipid levels in plasma of HD mice compared to WT mice, that were treated with ZFP-dDBD (**Fig. 5A**). Treatment of mice with ZFP-HTT corrected the total lipid levels in HD mice to WT levels. This is in contrast to the loss in total lipid levels in caudate-putamen in HD versus WT mice treated with control vector. Of note, inverse changes in total lipids were measured in plasma (increased, Fig. 5A) compared to caudate-putamen (decreased, Fig. 4B) and treatment status, suggesting a relationship between this peripheral biofluid and brain.

**Figure 5:**
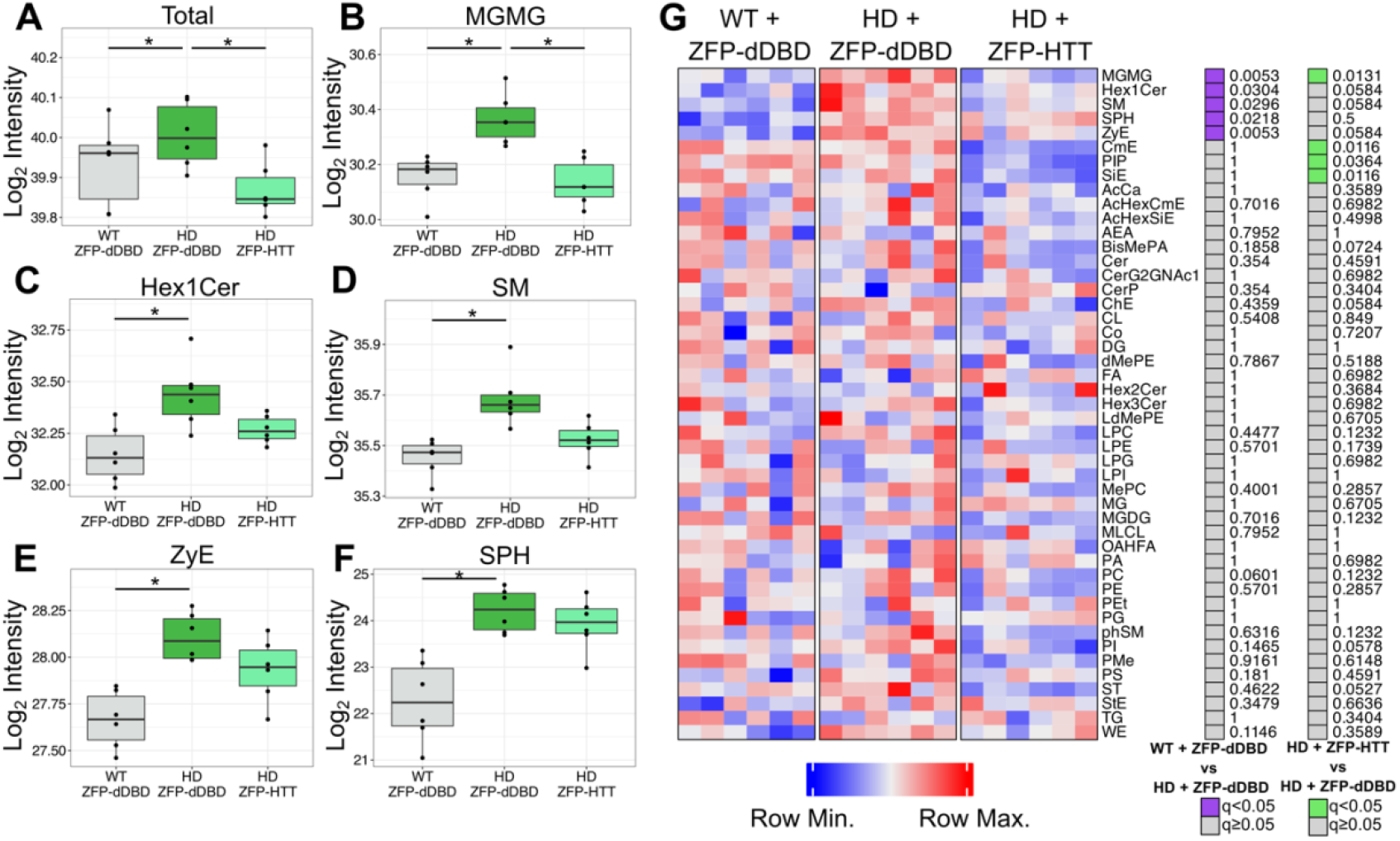
Protection of total lipid levels and subclass levels via intrastriatal lowering of mutant HTT in the plasma. **(A**) Total lipid levels are increased 5-10% in HD mice treated with ZFP-dDBD compared to WT treated with ZFP-dDBD and HD mice treated with ZFP-HTT, indicating recovery by ZFP-HTT. **(B)** MGMG levels are increased in HD mice treated with ZFP-dDBD compared to WT mice treated with ZFP-dDBD (Tukey’s Multiple Comparison; q = 0.0053,) and HD mice treated with ZFP-HTT (Tukey’s Multiple Comparison; q= 0.0013). **(C-F)** Hex1Cer, SM, ZyE, SPH were increased in HD mice treated with ZFP-dDBD compared to WT mice treated with ZFP-dDBD, there was no significant difference between HD mice treated with ZFP-dDBD and HD mice treated with ZFP-HTT. (* q<0.05)**(G)** The heatmap depicts individual animals (columns) and lipid subclass intensities (rows). *Purple* boxes indicate significantly different levels between WT + ZFP-dDBD and zQ175DN + ZFP-dDBD. *Green* boxes indicate significantly different levels between zQ175DN + ZFP-HTT and zQ175DN + ZFP-dDBD. All stats show are two-way ANOVA with Tukey’s multiple comparison test with a two-stage linear step-up procedure of Benjamini, Krieger and Yekutieli, N=44 lipid subclasses, n=6 mice.

Treatment with ZFP-HTT prevented changes for lipid subclasses in plasma **(Fig. 5G)**. There were five lipid subclasses significantly higher in HD mice compared to WT mice each treated with ZFP-dDBD **(Fig. 5 B-F)**. Notably, levels of MGMG and SM were increased in plasma and moved the opposite direction in the caudate-putamen (down). Additionally, MGMG was significantly lower in HD mice treated with ZFP-HTT compared to those treated with ZFP-dDBD, indicating MGMG subclass in plasma is an effective readout for therapy-responsive changes. Three of the other lipid subclasses: Hex1Cer, SM, and ZyE, were trending towards recovery.

Intrastriatal treatment with ZFP-HTT prevented many genotype-dependent changes in individual lipid ions in the plasma (**Supplemental File 1**). There were 70 significantly altered lipid ions between WT and HD mice treated with ZFP-dDBD, including species of PC. Of those, 56% (39/70) were protected by the ZFP-HTT treatment (**Table 1)**.

Overall, these results suggest that monitoring plasma was an effective method to evaluate the protective effects of intrastriatal injection of AAV-ZFP-HTT.

### AAV9-ZFP-HTT blocked somatic instability zQ175DN mice

Others have reported that ZFP-HTT can block SI in mice (29). We found there was a statistically significant decrease in SI index of 64% of the mutant HTT allele in HD mice treated with ZFP-HTT (2.59 ± 1.42) compared to ZFP-dDBD (7.33±0.877), (Unpaired t-test p = 0.014). This demonstrates that the ZFP-HTT can strongly inhibit CAG repeat expansion in the caudate-putamen of zQ175DN mice.

## Discussion

In this study we found numerous changes in lipid species between WT and zQ175DN HD mice at six months of age, and preservation of levels for most lipids with intrastriatal injection of a lowering therapy. Lipid changes could be measured in caudate-putamen and complimentary changes could be detected in peripheral biofluid (plasma). We also detected a misfolded species of mHTT resistant to lowering present in the crude nuclear fraction, reproducing earlier results in a different mouse HD model. Our results show that LC-MS offers a quantitative way to track biochemical changes in lipids reflective of neurodegeneration in preclinical models.

Treatment with AAV9-ZFP-HTT resulting in mHTT lowering was highly effective at preventing total lipid changes in caudate-putamen of zQ175DN mice. Thus, while complete reversal of all pathologies, including transcriptional changes, may require either very early mHTT lowering to prevent accrual of mHTT, or actively targeting accumulated mHTT in nuclear compartments, lowering mHTT with AAV9-ZFP-HTT still awarded significant benefits as measured by lipid changes.

Many of the lipid changes found in zQ175DN mouse brain in this study were like those found in synaptic fractions in zQ175/Q7 mice at six months of age in our previous study (23). Lipid changes in the LacQ140 mice, which have a shorter CAG repeat and a more prolonged disease process, were slightly different and changed with age, but also showed a significant loss of total lipids (at 9 months) with heavy losses of lipids important for white matter. Here, we also saw loss of lipids important for white matter health (SM, Hex2Cer, sulfatides), which can be lost with neurodegeneration and aging (30, 31), but lipid profiles were mostly unique to each animal model. The identity of lipids changed in HD mice (measured by us) and measured in HD autopsy tissue (32) are consistent with altered white matter, which reflects processes observed in HD patients and is likely to affect nerve conduction (reviewed by (33)).

Genotype- and treatment-dependent changes in lipids were detectable in mouse plasma. These results suggest a relationship between lipid levels in brain and plasma. Lipid alterations have been detected in plasma of HD patients, especially in the PC and free fatty acids (26-28). We measured changes in PC in HD mouse plasma, although the exact species of PC were different than those measured from HD patients. This could be due to species differences (human versus mouse) or extraction methods. We measured many more changes than prior studies using human plasma, which may be due to improved lipid annotation and increased sensitivity of instrumentation.

Lipid changes caused by mHTT may hinder neuronal function or alter white matter maintenance. mHTT immunoreactivity can be detected with anti-HTT antibodies PHP1 and PHP2 in myelin in zQ175/Q7 mice (34) positioning mHTT to contribute to pathology through post-transcriptional mechanisms such as interfering with membrane-associated protein complexes or changing lipid bilayer organization through direct lipid-mHTT interactions (35-37). The impact of mHTT in these compartments may be overcome by mHTT lowering. The misfolded species of mHTT resistant to lowering which we detected may also impair transcription including lipid metabolizing enzymes.

Carroll and colleagues (29) reported that binding of ZFP-HTT to the CAG on the HTT allele may block somatic instability providing additional protection from disease in addition to that afforded by mHTT protein lowering. A CAG repeat of 175 is thought to already exceed the length needed to induce most phenotypes including transcriptional phenotypes without further expansion (38). Even so, in addition to lowering mHTT levels, impeding somatic instability with ZFP-HTT may be one mechanism that contributes to the protection of lipid levels in our study.

In conclusion, assessment of lipid levels using a quantitative method revealed sweeping positive effects of a lowering therapeutic targeting mutant HTT transcription in a preclinical mouse model of HD.

## Acknowledgements

We thank David Howland and Deanna Marchionini at CHDI Foundation for helpful advice and Jonathon Asara at the Beth Israel Deaconess Medical Center Mass Spectrometry Core for help with LC-MS and lipid annotation.

## Supplementary Data

File 1. Excel file containing LC-MS results for all individual lipids % intensities p values, q values. Statistical analysis performed by ANOVA and posthoc Tukey’s test with 5% FDR correction.

## Supplementary Methods

Additional details of the methods and materials.

**Supplemental Figure 1:**
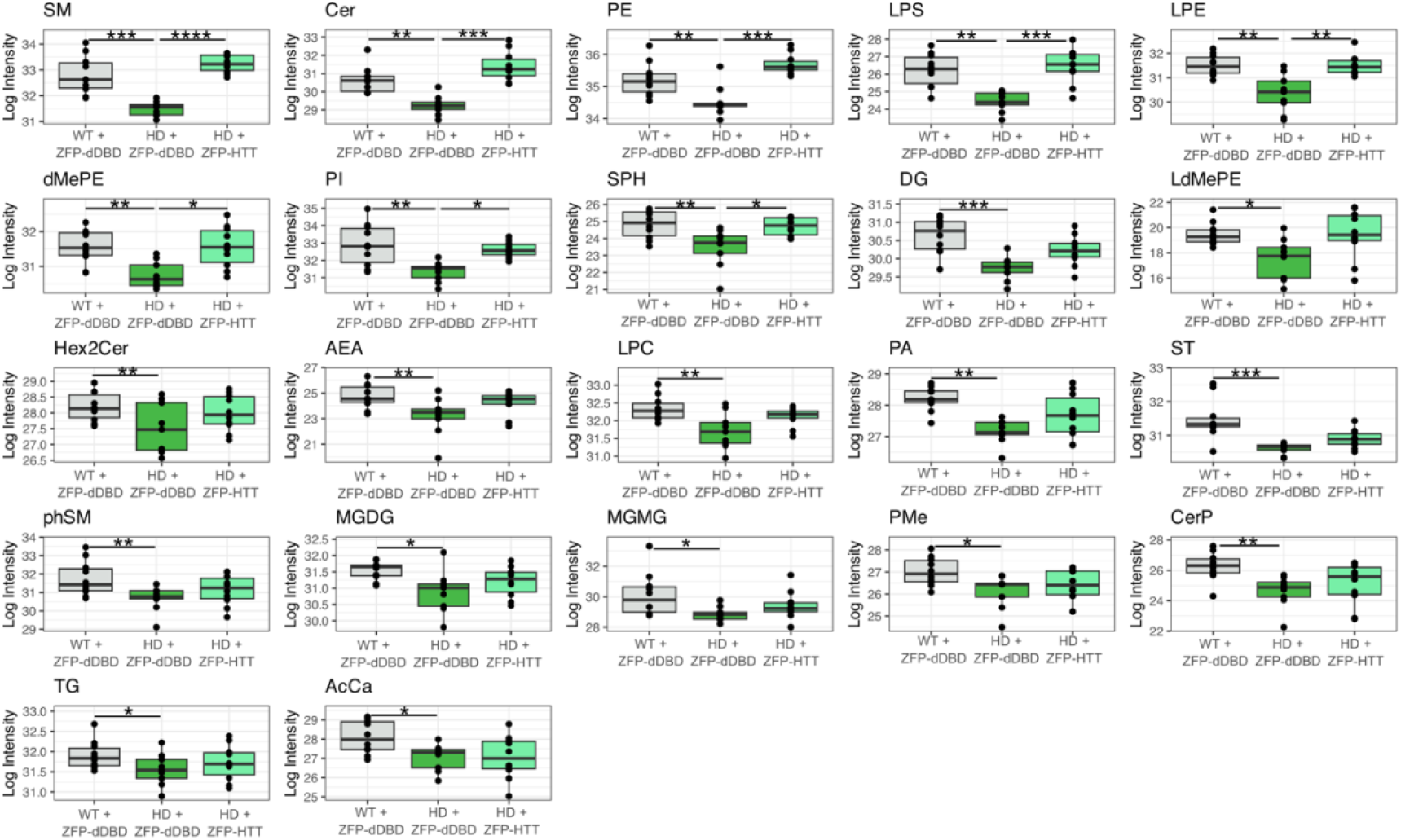
Significantly altered lipid subclasses in caudate-putamen of 6 month mice. Graphs shown mean and interquartile ranges of significantly altered lipid subclasses. All data was analyzed via Two-Way ANOVA with Tukey’s multiple comparison test and corrected with two-stage linear step-up procedure of Benjamini, Krieger and Yekutieli, (N=42 lipid subclasses, n=10 mice; ^*^q<0.05, ^**^q<0.01, ^***^q<0.001). SM, Cer, PE, LPS, LPE, dMePE, PI, SPH were corrected with treatment of ZFP-HTT. DG, LdMePE, Hex2Cer, AEA, LPC, PA, ST, phSM, MGDG, MGMG, PMe, CerP, TG, and AcCa, were not corrected with treatment of ZFP-HTT.

## Notes

### Competing Interest Statement

The authors have declared no competing interest.

### Summary of Updates

To accommodate new data concerning lipid changes Figure 5/Table 1 and somatic instability.

