## Supplemental Methods for "Intrastriatal delivery of a zinc finger protein targeting the mutant HTT gene by AAV9 obviates lipid phenotypes in brain and plasma of zQ175DN HD mice"

*Lipid Subclass Abbreviations*

| **Class** | **Subclass** | **Abbreviation** |
| --- | --- | --- |
|  | Acyl Carnitine | AcCa |
|  | N-acyl Ethanolamine | AEA |
| Fatty Acyl | Fatty Acid | FA |
|  | (O-acyl) -1- hydroxy fatty acid | OAHFA |
|  | Wax Ester | WE |
|  | Diacylglyceride | DG |
|  | Monoacylglyceride | MG |
| Glycerolipid | Monogalactosyl-diacylglycerol | MGDG |
|  | Monogalactosyl-monoacylglycerol | MGMG |
|  | Triacylglyceride | TG |
|  | Bis-methyl phosphatidic acid | BisMePA |
|  | Cardiolipin | CL |
|  | Dimethyl phosphatidylethanolamine | dMePE |
|  | Lysodimethyl phosphatidylethanolamineÍ | LdMePE |
|  | Lysophosphatidylcholine | LPC |
|  | Lysophosphatidylethanolamine | LPE |
|  | Lysophosphatidylglycerol | LPG |
|  | Lysophosphatidylinositol | LPI |
| Glycerophospholipid | Lysophosphatidylserine | LPS |
|  | Methyl-Phosphatidyl Choline | MePC |
|  | Mono-lysocardiolipin | MLCL |
|  | Phosphatidic acid | PA |
|  | Phosphatidylcholine | PC |
|  | Phosphatidylethanolamine | PE |
|  | Phosphatidylethanol | PEt |
|  | Phosphatidylglycerol | PG |
|  | Phosphatidylinositol | PI |
|  | Phosphatidylinositol-monophosphate | PIP |
|  | Phosphatidylmethanol | PMe |
|  | Phosphatidylserine | PS |
| Prenol | Coenzyme | Co |
|  | Ceramides | Cer |
|  | Simple Glc series | CerG2GNAc1 |
| Sphingolipid | Ceramide Phosphate | CerP |
|  | Ganglioside | GD1a |
|  | Ganglioside | GD1b |

|  | Hexosylceramide | Hex1Cer |
| --- | --- | --- |
|  | Dihexosylceramide | Hex2Cer |
| Sphingolipids (cont.) | Trihexosylceramide | Hex3Cer |
|  | Phyosphingomyelin | phSM |
|  | Sphingomyelin | SM |
|  | Sphingosine | SPH |
|  | Sulfatide | ST |
|  | AcylGlcCholesterol ester | AcHexChE |
|  | Cholesterol Ester | ChE |
|  | Zymosterol Ester | ZyE |
| Sterol | AcylGlcCamposterol Ester | AcHexCmE |
|  | Camposterol Ester | CmE |
|  | Sitosterol Ester | SiE |
|  | Sigamasterol Ester | StE |
|  | AcylGlcSitosterol Ester | AcHexSiE |

*AAV subcloning & packaging*

The zinc finger protein (ZFP) cDNA constructs, originally created by Sangamo, were engineered to bind to the *HTT* gene with specificity for expanded CAGs fused to the repressor domain of the KRAB protein (ZFP-HTT) (1) and were provided by CHDI Foundation. This fusion protein has been shown to selectively repress transcription from the mutant *HTT* allele containing high CAG repeats including the HTT1a exon1-intron1 readthrough transcript (1). The cDNA encoding construct “30645” specific to mutant HTT (ZFP-HTT) was fused in frame with the KRAB repressor domain (Construct D from Zeitler B. Nature Med 2019) and subcloned into the multiple cloning site of the self-complimentary adeno-associated virus 9 (AAV9) vector behind a chick beta-actin promotor **(Fig. 1A)**. The DNA binding domain (DBD) from this construct was deleted to serve as a control (ZFP-dDBD) (described by Zeitler B. Nature Med 2019) and was also subcloned into the same viral vector, and were transfected into HEK293T cells for AAV viral production as described (39, 40).

*Mouse Intracranial Injection Surgery and Tissue Collection*

The knock-in zQ175/Q7 mouse model was isolated as a natural variant in progeny of the Q140 knock-in mouse model and expanded (41, 42); the model was created by homologous recombination of mouse exon1 with human exon1 containing an expanded CAG expansion in the HTT allele. The neo cassette which suppress expression of the mutant HTT allele (43) was deleted to create zQ175/Q7 DN mice which express mHTT at endogenous levels. At eight weeks of age, mice were anesthetized with 284 mg/kg tribromoethanol (Avertin) and placed into a small animal stereotaxic apparatus (ASI Instruments and WPI Instruments). Microinjector pumps were fastened to the stereotaxic and set to deliver 2.5µL bilaterally (5µL total) of AAV9 virus (1 x 10^13^ PFU) at a rate of 500 nL/min. Our previous studies show that this dose results in near complete coverage of the caudate-putamen in mice. The surgical position was set using the bregma as the zero position for guidance, then measured ± 2.0 mm medial-lateral and +1.0 mm anterior-posterior (AP), and finally lowered -3.0 mm dorsal-ventral (DV) into the striatum (bilateral injections). At 6 months of age, Q7/Q7 WT or zQ175DN HD were deeply anesthetized with tribromoethanol (Avertin) before cervical dislocation. Whole brains were removed and frozen fresh at -80°C until use. Plasma was collected transcardially at the time of anesthetization. Samples were visually inspected for hemolysis. Plasma samples with the presence of hemolysis were excluded from further analysis.

*Tissue fractionation, SDS-PAGE, western blot analysis, capillary gel electrophoresis and filter trap assay.*

Tissue from freshly sacrificed mice (both striata) was processed as previously reported (23). Protein concentration was determined using the Bradford method (BioRad kit). Equal amounts of protein (10 μg) were separated by SDS-PAGE using 3-8% Tris acetate gels (Criterion 26 wells, BioRad). Crude homogenates were transferred to nitrocellulose using the Transblot Turbo and P1 samples were transferred to PVDF using a wet transfer overnight at 35V (22). Antibodies and dilutions are as follows: Anti-HTT Ab1 (aa1-17, (44), 1995, 1;2000, rabbit), Anti-HTT S830 (Generous gift of Dr. Gillian Bates, 1:6000, sheep), Anti-HTT MW8 (Developmental Studies Hybridoma Bank, University of Iowa, 1:500, mouse), Vinculin (MilliporeSigma, #V9131, 1:2000, mouse), GAPDH (MilliporeSigma, #MAB374, 1:10000, mouse). Secondary antibodies: Jackson ImmunoResearch1:2500 for rabbit, 1:5000 for mouse and sheep. For capillary gel electrophoresis the automated simple western system (Wes, ProteinSimple) was utilized to analyze equal amounts of protein (0.6 μg) with anti-polyQ antibody MW1 at a 1:50 dilution and Vinculin loading control at a 1:5000 dilution following manufacturer’s standard protocol for the 66-440kD assay kit. Aggregation assay was performed as previously reported (13, 45). Briefly, 40 μg P1 fraction was diluted to a final concentration of 2% SDS and vacuumed through a cellulose acetate membrane pre-equilibrated in 2% SDS using a dot blot apparatus (BioRad). Wells were washed with 2% SDS and the membrane was removed from the apparatus, washed in TBST then processed as for western blot as described above using anti-HTT antibodies S830 and MW8.

*Pixel intensity quantification*

Using ImageJ software, the area and average signal intensity of bands on western blot were determined by manually circling each band and multiplying to obtain the total signal intensity. For the P1 smear, the signal above the full length HTT band was measured. The total signal intensity for each protein was normalized to the vinculin or GAPDH loading controls. For the filter trap assay, the average signal intensity inside an equal size circle corresponding to the size of the well minus the background signal from an empty well was determined for each sample. Statistical analyses were unpaired t-test for mHTT levels by western blot and capillary electrophoresis and One-way ANOVA with Tukey’s multiple comparison test for all other proteins and filter trap assay.

*Lipid Extraction, LC-MS, and Annotations*

Briefly, 80 μl of crude homogenate from dissected mouse striatum in sucrose-HEPES buffer or 90 ul of plasma, were transferred to 7 ml glass scintillation vials containing 750 μl of HPLC grade methanol prior to extraction with MBTE. Lipid extracts were resuspended in 35 μl of a 1:1 mixture of methanol and 2-propanol, and hand delivered to the Beth Israel Deaconess Mass Spectrometry Core. The lipid extracts were analyzed by positive/negative ion switching LC-MS/MS and annotated using LIPIDSearch as described (46). Annotations are classified by LIPID MAPS subclasses.

*Somatic Instability Index*

Genomic DNA was isolated from striatal P1 samples (Qiagen DNEasy Blood and Tissue kit, Cat. 69504) and purified (New England Biolabs Monarch PCR and DNA Cleanup Kit, Cat. T1130) and average CAG number in HTT was determined by PCR and capillary gel electrophoresis of PCR products at the MGH Genomics Core. The starting (inherited) CAG for each mouse was provided by Jackson laboratories from tail DNA. Somatic Instability Index was calculated as described, using a custom Python script. (47).
